# Spatiotemporal cerebral blood flow dynamics underlies emergence of the limbic-sensorimotor-association cortical gradient in human infancy

**DOI:** 10.1101/2024.04.10.588784

**Authors:** Minhui Ouyang, John A. Detre, Jessica L. Hyland, Kay L. Sindabizera, Emily S Kuschner, J. Christopher Edgar, Yun Peng, Hao Huang

**Affiliations:** Department of Radiology, Children’s Hospital of Philadelphia, 3401 Civic Center Boulevard, Philadelphia, PA, 19104, United States; Department of Radiology, Perelman School of Medicine, University of Pennsylvania, 3400 Spruce Street, Philadelphia, PA 19104, United States; Department of Neurology, Perelman School of Medicine, University of Pennsylvania, 3400 Spruce Street, Philadelphia, PA 19104, United States; Department of Psychiatry, Perelman School of Medicine, University of Pennsylvania, 3400 Spruce Street, Philadelphia, PA 19104, United States; Department of Radiology, Beijing Children’s Hospital, Capital Medical University, Beijing, 100045, China

**Keywords:** cerebral blood flow, brain development, limbic-sensorimotor-association cortical gradient, infant, behavior, neurodevelopmental outcome, arterial-spin-labeled MRI

## Abstract

Infant cerebral blood flow (CBF) delivers nutrients and oxygen to fulfill brain energy consumption requirements for the fastest period of postnatal brain development across lifespan. However, organizing principle of whole-brain CBF dynamics during infancy remains obscure. Leveraging a unique cohort of 100+ infants with high-resolution arterial spin labeled MRI, we found the emergence of the cortical hierarchy revealed by highest-resolution infant CBF maps available to date. Infant CBF across cortical regions increased in a biphasic pattern with initial rapid and sequentially slower rate, with break-point ages increasing along the limbic-sensorimotor-association cortical gradient. Increases in CBF in sensorimotor cortices were associated with enhanced language and motor skills, and frontoparietal association cortices for cognitive skills. The study discovered emergence of the hierarchical limbic-sensorimotor-association cortical gradient in infancy, and offers standardized reference of infant brain CBF and insight into the physiological basis of cortical specialization and real-world infant developmental functioning.

## Introduction

Although human brain growth continues into adulthood, it is most dramatic in infancy, doubling in size during the first year and to almost adult size by age 5 ^1,2^. The brain is one of the most metabolically expensive organs in the body; despite representing only about 2% of total body weight, the adult human brain consumes nearly 20% of the body’s basal metabolism^3^. However, the infant brain uses a strikingly larger fraction of the total body energy consumption, exceeding 40% ^4,5^. Such high energy consumption is vital for supporting intensive maturational processes, including synaptogenesis, glial proliferation, and myelination ^6^. Cerebral blood flow (CBF) delivers both oxygen and glucose, the principal brain substrates for energy production, to the brain. Because of the tight coupling between regional CBF (rCBF) and local cerebral metabolism ^7,8^, rCBF can be used as a surrogate measure of regional brain metabolism. The “brain growth spurt” during infancy ^9^ is associated with enhanced vulnerability to disorders ^10^. If blood flow is interrupted or inadequately supplied, the brain is more susceptible to disorders such as ischemic stroke and hypoxic ischemic brain injury. Infancy is also a crucial period of behavioral development, with proper rCBF needed to support the emergence of cognitive abilities and behaviors. Accordingly, characterization of normative infant rCBF trajectories provides an opportunity to explore how brain regions use energy differentially in support of their functional specializations in health and disease, as well as the physiological basis for the progression of real-world developmental functioning. However, information on how cortical rCBF topography evolves throughout infancy and its associations with behavioral development is surprisingly lacking.

Differential cortical development ^11,12^ characterized by certain cortical gradient is one of the major hierarchical organizing principles of the human brain. Although the sensorimotor-association gradient has been well characterized in adolescents and adults ^13^, such a pattern cannot be completely discerned at birth. During infancy, cortical gradient may exhibit a slightly different pattern than in later developmental stages. The gradient could involve other components such as the limbic system, given the phylogenetically old nature of limbic functions and babies’ tendency to behave in a limbically motivated manner. Tremendous developmental brain changes occurring throughout infancy are critical for the emergence of cortical gradients. Neuroimaging derived measures of infant brain ^14^ such as cortical morphology ^e.g. 15,16^, white matter microstructure e.g. ^2,17–19^, and functional connectivity ^e.g. 20–22^ suggest a hierarchical sequence of cortical development in which limbic and primary areas mature before association cortices. These developmental brain changes demand rapid increases in metabolic substrate. Previous fluorodeoxyglucose positron emission tomography (FDG-PET) studies suggest a spatially heterogenous courses of metabolic maturation across infant cortex, with increases in local cerebral glucose utilization observed in temporal and primary visual cortices at 2-3 months of life, and then in the frontal cortex at 6-12 months ^23,24^. The dynamic course of local brain metabolism revealed in these studies provides grounds for anticipating a spatiotemporally heterogenous pattern of rCBF trajectory during infancy. We thus hypothesized that the infant rCBF trajectory proceeds in a hierarchical manner, beginning with limbic and primary cortices at the bottom and gradually transitioning to heteromodal association areas toward the top of the hierarchical levels.

To date, there has been little, if any, direct rCBF-based physiological evidence supporting the emergence of limbic-sensorimotor-association cortical gradient. Complicated by both neural and non-neural components ^25,26^, the mechanisms of fMRI blood-oxygen-level-dependent (BOLD) signal remain elusive and resultantly fMRI-based mapping of cortical hierarchy likely reflect measures of neural activities mixed with other factors such as vasomotor activity. Complementary to the valuable fMRI BOLD studies, our study exclusively based on fundamental physiological CBF measurements that directly and quantitively reflect regional resting state neural activity, could offer solid and refreshing insight into the emergence of the limbic-sensorimotor-association cortical gradient. Arterial spin labeled (ASL) MRI uses magnetically labeled arterial blood water protons as an endogenous tracer for reliably estimating rCBF ^27^. As compared to FDG-PET, which requires the intravenous administration of a radioactive tracer, ASL MRI is completely noninvasive and hence well suited for use in healthy infants and children ^e.g.28–32^. Collectively, there is still a knowledge gap in not only mapping infant rCBF to regional precision but also depicting the organizing principle of rCBF dynamics that underpin cortical gradient emergence during the critical developmental period from 0 to 2 years.

In this study, by acquiring the highest-resolution infant CBF maps available to date with densely sampled ages during infancy, we explored the organizing principle of spatiotemporal dynamics of rCBF in typically developmental infants with greater precision. The high-resolution spatiotemporal dynamics of rCBF were obtained by capitalizing on a state-of-the-art pseudo-continuous ASL (pCASL) imaging protocol with 3D multi-shot stack-of-spirals readout ^33,34^ and background suppression. Unlike previous studies in the setting of clinical suspicion of diseases, a typically developmental infant CBF study with such a scale of samples within 0-2 years is unprecedented. We found that the global CBF increased in a logarithmic fashion during infancy. The high-resolution, high-quality infant rCBF maps revealed physiological heterogeneity in spatial distribution as well as age-related increases. These findings were also replicated in a second cohort using high-resolution 2D pCASL imaging. By employing a data-driven clustering approach to identify topological regionalization of rCBF with a similar changing pattern, we further discovered three distinct clusters, including limbic, sensorimotor, and frontoparietal areas that reflect the limbic-sensorimotor-association cortical gradient. Lastly, we found that rCBF changes were associated with improved real-world developmental functioning across multiple domains. Together, these findings suggest that the spatiotemporal infant rCBF dynamics underlie the emergence of a hierarchical limbic-sensorimotor-association gradient and contribute to the development of behavior in infancy, setting the stage for cortical function specialization.

## Results

### Infant participant profile

A total of 134 infants from two cohorts (N=78 for cohort-1 and N=56 for cohort-2; see Participants in Methods and Supplementary Table 1) participated this study. Briefly, cohort-1 is from an ongoing study at the Children’s Hospital of Philadelphia. This study investigates normal brain development during infancy. All infants from cohort-1 are typically developing children and recruited completely for research purposes. Cohort-2 is from Beijing Children’s Hospital. Each infant in cohort-2 had their clinical history thoroughly reviewed to rule out developmental abnormalities, and no brain abnormalities were detected through reading of an experienced pediatric radiologist (YP) of these infants’ MRI scans. It should be noted that almost all results (Figs 2-6 and almost all supplemental Figures) were obtained from one cohort (cohort-1) of infants in one scanning location.

### Logarithmic increase of global CBF during human infancy revealed by datasets of 100+ infants with densely sampled ages

As a first step, we established the infant developmental curve of global CBF with a relatively large datasets from 119 infants (60M/59F; Age range of 1.4-28 months with mean/standard deviation 12.5 ± 7.1 months) who had completed phase-contrast (PC) perfusion MRI in both cohorts. Global CBF for each infant was measured by PC perfusion MRI, a reliable global CBF measurement technique ^35^ that encodes the velocity of flowing spins as phase changes in an MRI image. By measuring the flow flux at the four feeding arteries of the brain (Supplementary Fig. 1A), including bilateral internal carotid arteries and vertebral arteries, global CBF can be noninvasively measured. We investigated the effects of age, sex, and cohort on global CBF using a large infant database from these 119 infants (see Participants in Methods). We found that global CBF increased in logarithmic manner during infancy (*r* = 0.823, *P* < 2×10^-16^; Fig. 1), after testing and quantifying the relative goodness-of-fit among different models (including linear, logarithmic, exponential, Poisson, and quadratic polynomial) for fitting the developmental curve (Supplementary Fig. 2 and Supplementary Table 2). Sex had no effect on global CBF trajectory, nor did cohort have any detectable effect (Supplementary Table 2). Segmented regression analysis revealed two distinct phases of global CBF increases during infancy (*F* (2,117) =22.23, *P* < 6.88×10^-9^; Supplementary Table 4), with a similar breakpoint as that of the total daily energy expenditure growth ^36^ (see Discussion). Global CBF rose rapidly at 3.52 ± 0.45 ml/100g/min per month in the first phase from birth to a break point at 10.75 months of age (95% CI - confidence interval: 8.70, 12.81), followed by sequentially slower increases at a rate of 0.43 ± 0.23 ml/100g/min per month in the second phase. These findings highlight significant physiological changes that occur during infant brain development and offer a normative reference of global CBF during infancy.

**Figure 1.**
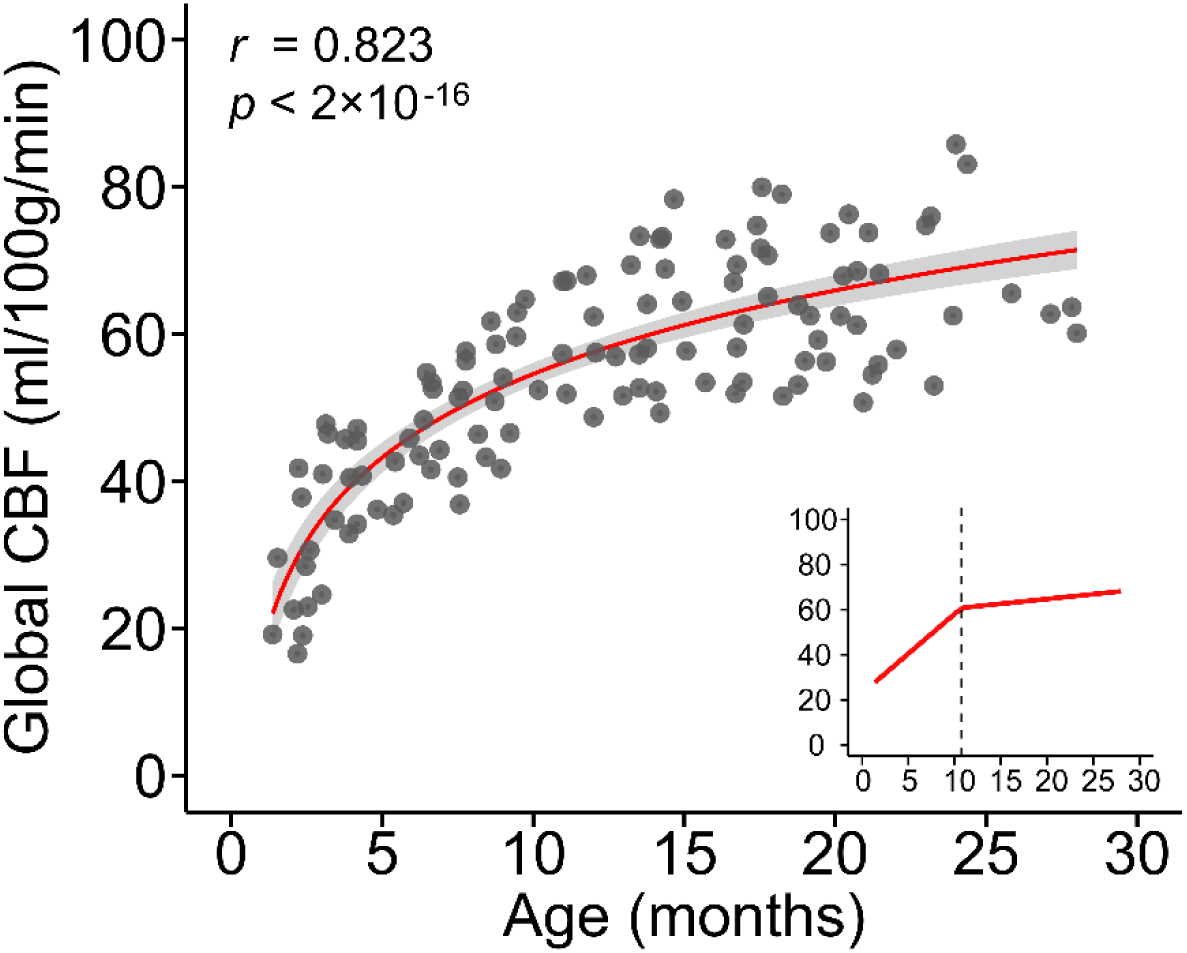
Developmental curve of global cerebral blood flow (CBF) throughout infancy. Global CBF increases in a logarithmic manner during infancy (*r* = 0.823, *P* < 2×10^-16^). The red line in the main panel indicates the best linear fit of global CBF = 16.38×log(age)+16.85 (Supplementary Fig. 2 and Supplementary Table 2), and the shaded envelope denotes the 95% confidence interval. Data points represent global CBF measured with phase-contrast MRI of each infant (*N* = 119). In the bottom-right panel, segmented regression analysis indicates a biphasic pattern of global CBF increase (red line) with a break point at 10.75 months. The fitted bilinear regression line was generated with segmented regression analysis (See Methods section).

### Regional physiological variability at finer detail across infant brain revealed by high-resolution rCBF maps based on arterial spin labeled (ASL) perfusion MRI

Having established the normative developmental trajectory of global CBF, we then examined how regional CBF topography evolves with age during infant development. Due to the small size of infant brains, substantially higher spatial resolution usually at the cost of signal-to-noise ratio (SNR) ^2^ of rCBF images is required to achieve comparable “anatomic” details to the adult brain. To achieve high spatial resolution and high SNR, we leveraged an advanced 3D multi-shot, stack-of-spiral pCASL MRI sequence ^33,34^ and developed an infant-dedicated protocol to acquire *in vivo* ASL images at 2.5mm isotropic resolution. In contrast to resolutions as low as 4-10mm conventionally used in infant and pediatric populations, the resulting datasets acquired at the single Philadelphia site (*N* = 76, 32M/44F; Supplementary Table 3) provided the highest resolution ASL data to date for infants, and revealed heterogeneous spatial distribution patterns (Fig. 2 and Supplementary Fig. 3) that could not be observed previously. Higher resolution and SNR with the cutting-edge pCASL ^33, 34^ can be clearly observed when compared with conventional protocol (Supplementary Fig. 4). The dense acquisition of thin axial slices in our infant-dedicated pCASL protocol facilitates more precise estimation of regional CBF. These highest resolution infant CBF maps available to date capture around 3 times of cortical voxels captured in the conventional rCBF maps (Supplementary Figure 4), revealing the unprecedented details of the whole-brain physiological dynamic patterns in infants. The rCBF findings shown in Figures 2-6 are from 76 typically developing infants in cohort-1 only. To ensure reliability of the datasets, we examined the reproducibility of rCBF maps by scanning an infant twice using the same scanner and identical pCASL protocol (Supplementary Fig 1B). Consistent with literature on high test-retest reliability of ASL MRI^37^, the intraclass correlation coefficient of two scans in this study was 0.869 (95% CI: 0.861,0.876) (Supplementary Fig. 1C), indicating excellent reliability of rCBF maps obtained with our pCASL protocol.

The resulting higher resolution and effective SNR of infant rCBF maps enable more precise localization and provides finer details for physiological variability in the infant brains than prior work^29–31^. As demonstrated by rCBF maps from three representative infants aged 2, 8 and 17 months in Supplementary Fig. 3, the densely acquired axial slices of the infant rCBF maps reveal a high gray and white matter contrast (Supplementary Fig. 3A-C). After mapping to the cortical surface, regional heterogeneity of rCBF across infant cortex can be clearly appreciated (Supplementary Fig. 3D-F). Specifically, we found that rCBF values in the 2-month-old infant brain are higher in primary cortex (sensorimotor, auditory, and visual) and lower in association cortex (frontal and parietal), whereas in the 17-month-old infant, higher rCBF values are more prominent in association cortex. We further calculated group-averaged rCBF maps at six milestone age groups (Fig. 2). Consistent with the spatial distributions observed in individual infants, we discovered that higher values in group-averaged rCBF maps are preferentially exhibited in primary sensory cortex in infants younger than 6 months (white arrows in Fig. 2) versus association cortex in infants older than 12 months. These observations demonstrate good spatial resolution and SNR of the infant rCBF maps to capture spatiotemporal variability for us to further tackle the emergence of cortical hierarchy from the physiological aspect.

**Figure 2.**
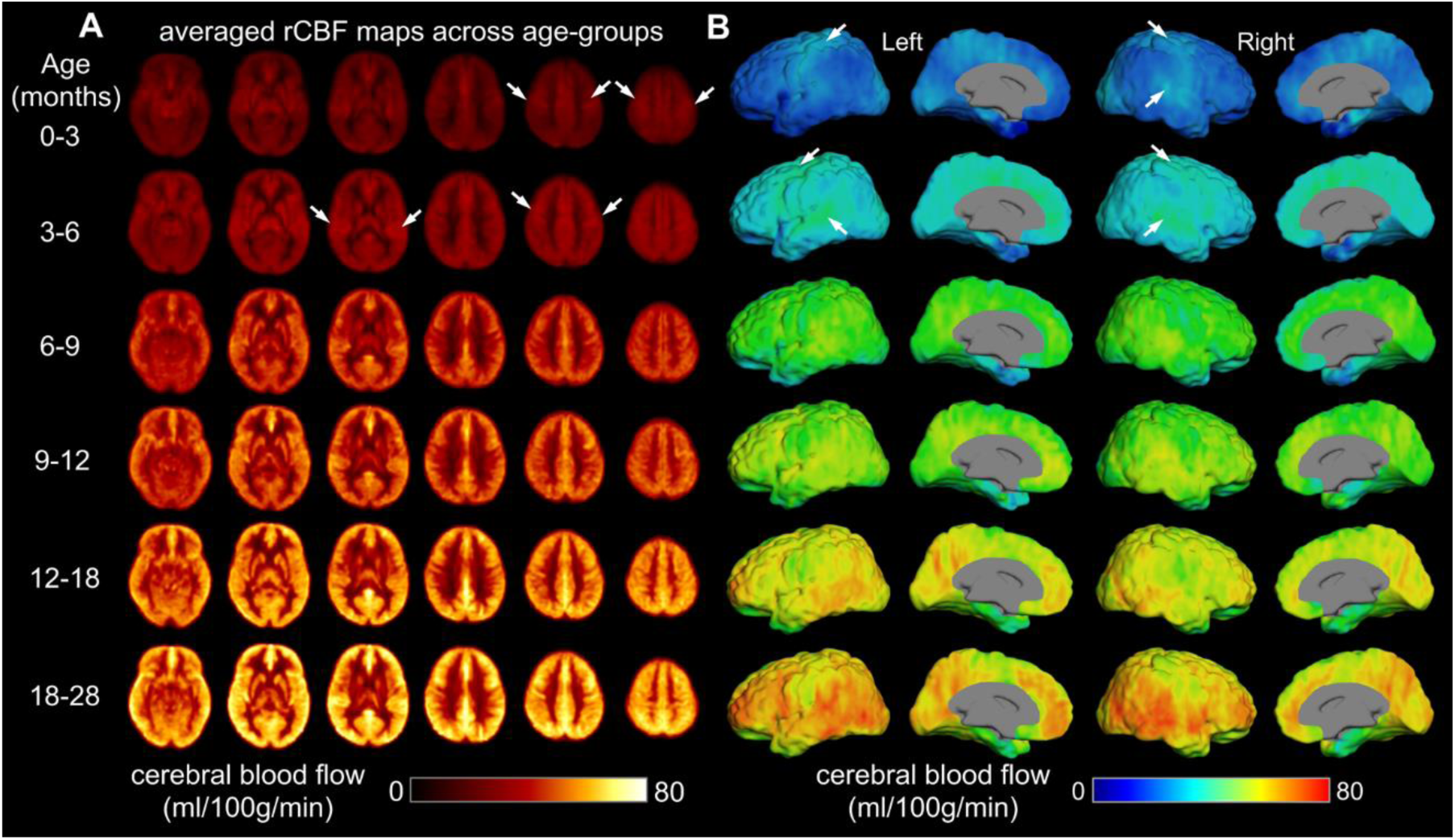
Precise physiological variability at finer detail across infant brain regions. Population-averaged regional cerebral blood flow (rCBF) maps in template space from infants age groups of 0-3, 3-6, 6-9, 9-12, 12-18 and 18-28 months. High-resolution (2.5x2.5x2.5mm^3^) rCBF maps were acquired with 3D multi-shot, stack-of-spirals pseudo-continuous arterial spin labeled (pCASL) perfusion MRI. **(A)** Six representative axial slices of averaged rCBF maps from inferior to superior are shown from the left to right for each age group. **(B)** Averaged rCBF maps are projected to the 3D reconstructed surface of a template brain and displayed in lateral and medial view of both hemispheres. White arrows indicate the high CBF values in the primary sensorimotor and auditory cortices at younger age groups.

### Nonuniform age-related regional CBF increases across heteromodal and unimodal cortex during infancy

Based on the empirical observation that higher rCBF values are located in the unimodal primary cortex of younger infants and in the heteromodal association cortex of older infants (Fig. 2 and Supplementary Fig. 3), we hypothesized that age-related changes in rCBF would vary heterogeneously across cortical regions. To test this hypothesis, we used the high-resolution infant rCBF dataset from cohort-1 (*N* = 76) and logarithmic models to examine how rCBF is associated with age during infant development, based on a logarithmic pattern for global CBF shown in Fig. 1. Sex and in-scanner motion were included as covariates in these models. After correcting for multiple comparisons with the Bonferroni method (Z ≥ 5.1, Bonferroni-corrected p value *P_bonf_* ≤ 0.05), we found that age-related increases in rCBF were nonuniform across the cortex (Fig. 3A). Specifically, the most significant age-related increases in rCBF were seen in areas of heteromodal association cortices, including prefrontal cortex, inferior parietal lobule, lateral temporal cortex, precuneus, and posterior cingulate cortex. Substantial rCBF increases were also observed in unimodal sensorimotor cortex, but to a lesser extent.

**Figure 3.**
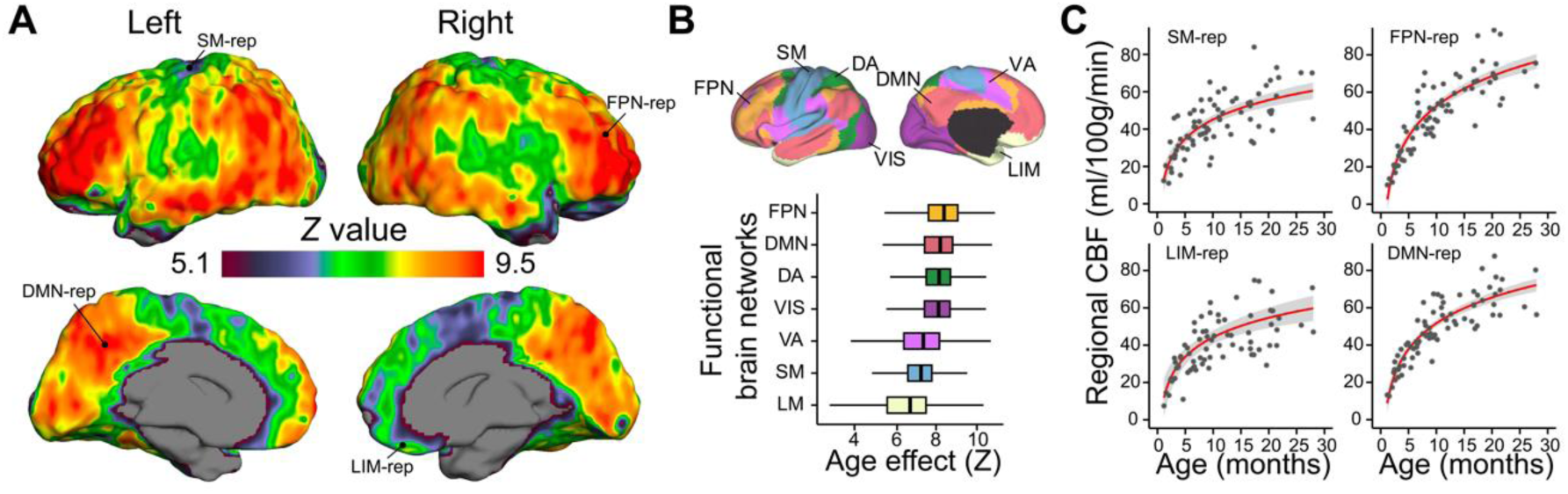
Nonuniform age-related increases of regional CBF during infancy. **(A)** Regional CBF increases in a logarithmic fashion across the cortex, most prominent in the heteromodal association cortex than unimodal cortex. Images thresholded at z > 5.1 (Bonferroni corrected p < 0.05). **(B)** Regional CBF increases varied heterogeneously by functional brain networks defined by Yeo et al., (2011) (top panel). The box plots in the bottom panel reflect the voxel-wise age effect of infant rCBF in seven functional networks ordered by median value. Regional CBF increases most significantly with age in the fronto-parietal and default-mode networks, and less so in the limbic and sensorimotor networks. DA: dorsal attention; DMN: default-mode network; FPN: fronto-parietal network; LIM: limbic; SM: sensorimotor; VA: ventral attention; VIS: visual. **(C)** The developmental curves of infant rCBF from representative voxels located in four brain networks: sensorimotor as SM-rep (*upper-left panel*), limbic as LIM-rep (*bottom-left panel*), fronto-parietal as FPN-rep (*upper-right panel*) and default-mode as DMN-rep (*bottom-right panel*). Voxel locations were indicated in (A). Data points in scatter plots represent rCBF measured with advanced pCASL for each infant. The red lines in the right panels indicate the best linear fit of rCBF, and the shaded envelope denotes the 95% confidence interval.

Our findings were further supported by a validation analysis of voxel-wise age effects using rCBF maps (*N* = 48) obtained from cohort-2 (Beijing site) ^30^. The age effects were highly reproducible in both infant rCBF datasets, showing similar spatial topographies (Supplementary Fig. 5A-B) and high between-dataset correspondence (*r* = 0.426, *P_perm_* < 10^-4^; Pearson correlation with permutation test; Supplementary Fig. 5C). The infant rCBF dataset from cohort-1 in Philadelphia (Fig. 3A and Supplementary Fig. 5A) tended to yield stronger age effects than the validation cohort from Beijing (cohort-2; Supplementary Fig. 5B), most likely due to differences in pCASL acquisition (voxel size of 2.74×2.74×5 mm^3^ for the Beijing cohort versus the higher resolution of 2.5×2.5×2.5 mm^3^ for the Philadelphia cohort). Additionally, infants in the Beijing cohort (cohort-2) were scanned due to clinical indications, although no brain abnormalities were detected in cohort-2 infants based on an experienced pediatric radiologist’s MRI reading. Infants in the Philadelphia cohort (cohort-1) were typically developmental ones recruited solely for research.

We further examined the age effects of regional CBF using both predefined functional networks from Yeo brain network atlas ^38^ (Fig. 3B-C) and a data-driven clustering approach (Fig. 4) with high-resolution rCBF dataset from the cohort-1. Notably, age effects were nonuniformly distributed across seven functional networks (Fig. 3B), with the largest increases in rCBF occurring in fronto-parietal and default-mode networks, and with modest increases in rCBF observed in sensorimotor and limbic networks. For instance, rCBF from the fronto-parietal and default-mode network regions demonstrated stronger age effects (FPN-rep: *β* = 24.08, *r* = 0.891, *P* < 2×10^-16^; DMN-rep: *β* = 20.21, *r* = 0.887, *P* < 2×10^-16^) than rCBF from sensorimotor and limbic network regions (SM-rep: *β* = 15.44, *r* = 0.757, *P* = 6.51×10^-14^; LIM-rep: *β* = 16.59, *r* = 0.632, *P* = 6×10^-9^) (Fig. 3C). We constructed rCBF maps from the developmental curves at various ages throughout infancy (Supplementary Figs. 6-7). The resulting trajectory was animated to create a quantitative time-lapse movie of infant rCBF dynamics (Supplementary Movie 1). Together, the spatiotemporal rCBF measures revealed a shifting pattern of distribution with its developmental gradient beginning in limbic and sensorimotor areas and spreading rostrally into prefrontal cortex, as well as dorsally and caudally into parietal and temporal cortices.

**Figure 4.**
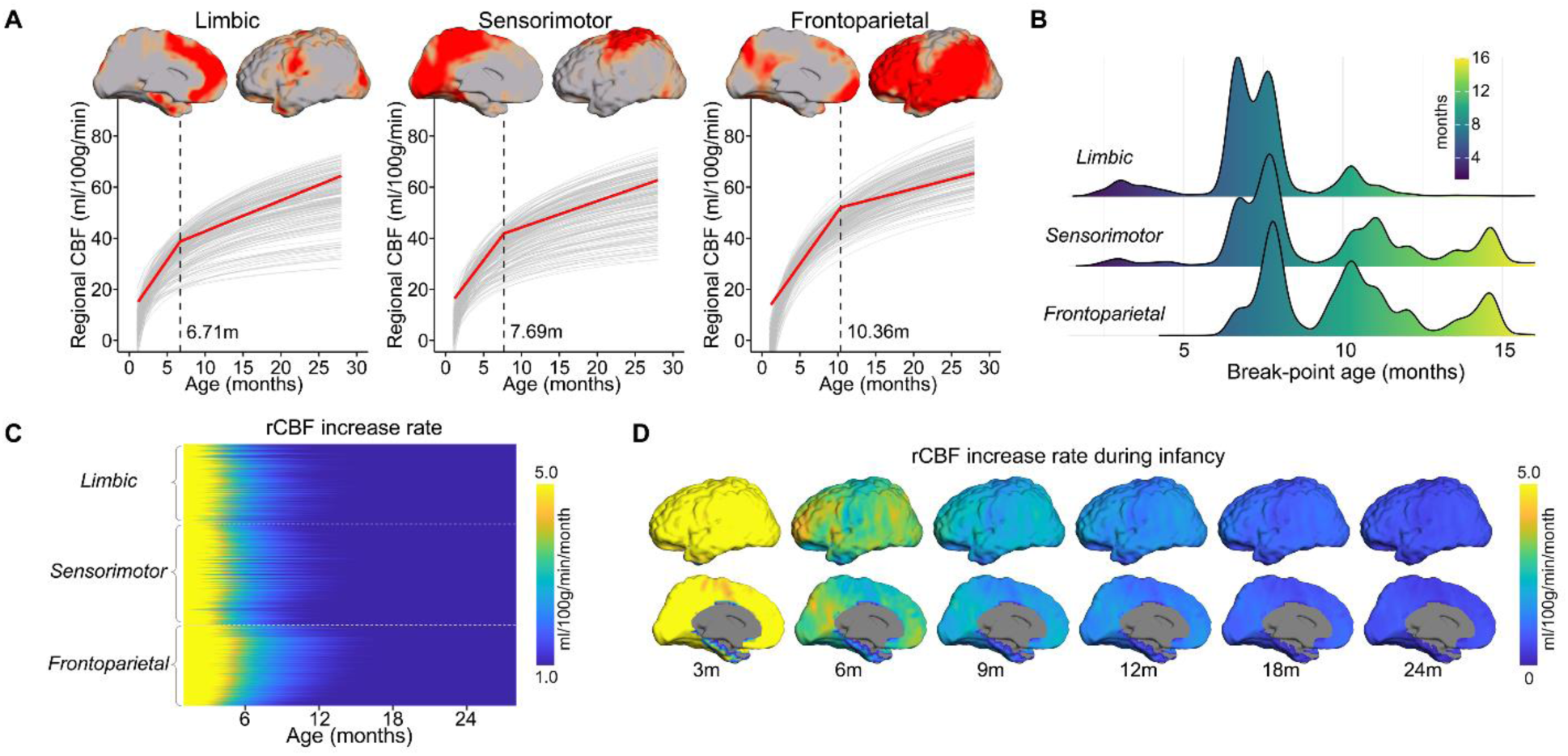
Infant rCBF increases according to a hierarchical limbic-sensorimotor-association gradient. **(A)** Cortical voxels were clustered into three groups (limbic, sensorimotor, and frontoparietal clusters) identified by non-negative matrix factorization (NMF) according to the pattern of rCBF increase over time. The respective locations of the three clusters on the cortical surface are displayed in lateral and medial views. Segmented regression analysis indicates a biphasic developmental pattern of averaged rCBF in each cluster (red line) overlaid on 200 randomly sampled voxel developmental curves from each cluster (gray thin lines). The identified break-point ages (black dashed lines) from segmented regression analysis varied across clusters and were provided at the bottom of each plot. **(B)** Histograms showed the profile of break-point age from cortical voxels within each cluster. **(C)** rCBF increase rate (ml/100g/min/month) across cortical voxels within each cluster. **(D)** Heterogeneous rCBF increase rate across cortex at milestone ages during infancy. Slower and faster rCBF increase rates are shown in cool and warm colors, respectively.

### Hierarchical limbic-sensorimotor-association gradient representing a general organizing principle of infant regional CBF dynamics

The findings above demonstrate notable areal heterogeneity in the age effect of rCBF, prompting the brain to be divided into clusters (e.g. groups of voxels), each with a more similar rCBF increase pattern than the rest of the brain. To provide a more nuanced understanding of the physiological dynamics during infancy, we performed data-driven clustering analysis of infant rCBF maps using non-negative matrix factorization (NMF)^39^ to identify distinct clusters. The NMF approach has been shown to identify brain networks robustly and accurately using neuroimaging data ^e.g., 11^. Only high-resolution rCBF maps from 76 TD infants from cohort-1 were included in these data-driven clustering analyses. We discovered three clusters along the hierarchical limbic-sensorimotor-association gradient with unique rCBF increase patterns (Fig. 4A and Supplementary Fig. 8). By comparing with the commonly used Yeo brain network atlas ^38^, voxels corresponding to each cluster primarily highlighted limbic, sensorimotor, and frontoparietal regions, reflecting limbic/sensorimotor-association hierarchy. To further examine the break-point ages between faster and slower rCBF increases, we conducted segmented regression analysis (elaborated in global CBF analysis) on averaged rCBF values from each cluster, resulting in similar biphasic patterns for all clusters (all *F* (2,74) ≥ 5.61, p ≤ 5.4x10^-3^). RCBF measures for all clusters increase faster in the first phase and then increase steadily but at a much slower rate in the second phase (Fig. 4A and Supplementary Table 4). The identified break points differed across clusters and exhibited a clear pattern along the hierarchical gradient. Notably, the break point was earliest for the limbic cluster at 6.71 months (95% CI: 4.09, 9.33), followed by the sensorimotor cluster at 7.69 months (95% CI: 5.63, 9.75), and then lastly the frontoparietal cluster at 10.36 months (95% CI: 8.21, 12.50). We also examined the break point profiles of all voxel-wise rCBF trajectories in the infant cortex. The resulting histograms (Fig. 4B) demonstrate that voxels in cortex from higher hierarchical levels (e.g., frontoparietal regions) have older break-point ages than voxels from lower hierarchical levels (e.g., sensorimotor and limbic regions). The robustness of our findings at the cluster and voxel-wise levels demonstrates that the biphasic pattern of rCBF increases varies along a cortical hierarchy, from limbic, and sensorimotor to frontoparietal association cortex.

Rates of regional infant rCBF increase calculated as the first derivative of fitted logarithmic curves of all cortical voxels further supports the limbic-sensorimotor-association gradient. We observed the highest rates of rCBF increase before 6 months, followed by dramatically slower increase rates after 6 months (Fig. 4C). Intriguingly, the duration of the rapid increase is longer in regions with higher hierarchical levels across three clusters (Fig. 4C). The rCBF increase rate maps at six milestone ages are demonstrated in Fig. 4D.

### Regionally specific rCBF increases supporting the emergence of infant behavior and real-world developmental functioning

With the NMF parcellation in Fig. 4A, we sought to determine where the rCBF increases support certain enhanced behavior during infancy by investigating the association between the rCBF and measures of infant real-world developmental functioning. Using the Bayley scales of infant and toddler development, we assessed infants’ behavior and functioning in the areas of motor, language, and cognition. We employed generalized additive models (GAMs)^40^ to model the association between all voxel-wise rCBF measures and infant neurodevelopmental outcomes, controlling for age, sex, in-scanner motion, and family socioeconomic status. Among the 76 infants who had high-resolution pCASL acquired in cohort-1, 49 of them have their Bayle scales and family socioeconomic status available and are included in the association analysis here. Cohort-2 is not included for the rCBF-behavior analyses. The results were clusters of voxels significantly correlated with Bayley domain scores (Fig. 5A-C). Specifically, the largest clusters of infant rCBF significantly correlated with motor (e.g. R.ANG, *t* = 3.670, *P_bonf_* = 0.007; Fig. 5A) and language skills (e.g. L.MOG/IOG, *t* = 3.703, *P_bonf_* = 0.006; Fig. 5B) were found in the sensorimotor/frontoparietal cortices. RCBF cluster located in the frontoparietal cortex (e.g. R.ANG, *t* = 3.624, *P_bonf_* = 0.007; Fig. 5C) significantly correlated with cognitive skill. The complete list of clusters is shown in Supplementary Table 5.

**Figure 5:**
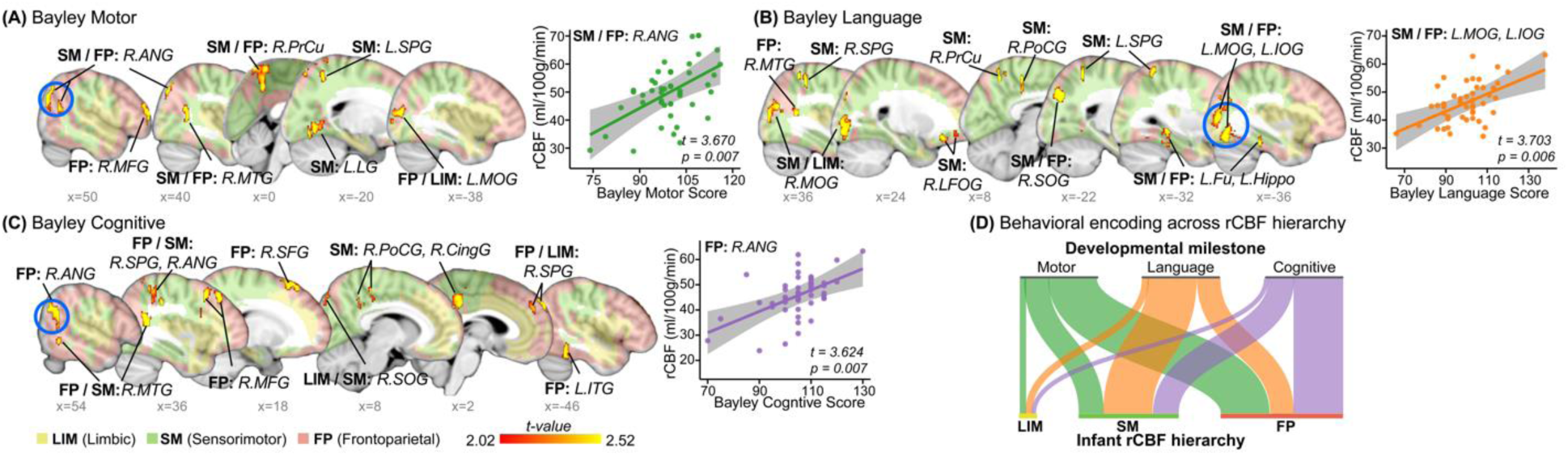
Regionally specific rCBF increases are associated with real-world developmental functioning in infants. Of all cortical regions examined, the regions in red-yellow showed significant positive associations (*p* _corrected_ < 0.05, cluster with *k* > 100 voxels, *t* > 2.02) between rCBF in infants and their behavior and developmental functioning quantified with Bayley scales of infant and toddler development across domains of **(A)** motor, **(B)** language, and **(C)** cognitive. The identified three clusters across infant rCBF hierarchy are shown in different colors in panel A-C, with limbic in light yellow, sensorimotor in light green and frontoparietal in pink. Scatter plots show the significant positive correlations between infant’s behavioral scores and the averaged rCBF values from the largest, blue circled clusters (Bonferroni corrected *p < 0.05*). The bold line indicates the best linear fit, and the shaded envelope denotes the 95% confidence interval. **(D)** River plot shows spatial distribution of voxels with significant association of each score across the identified rCBF hierarchy. Ribbons are normalized by the total number of voxels with significant associations in each behavioral score, shown in a different color. LIM, limbic; SM, sensorimotor; FP, frontoparietal; L/R, left/right hemisphere; ANG, angular gyrus; CingG, cingulate gyrus; Fu, fusiform gyrus; Hippo: hippocampus; IOG, inferior occipital gyrus; ITG, inferior temporal gyrus; LFOG, lateral fronto-orbital gyrus; LG, lingual gyrus; MFG, middle frontal gyrus; MFOG, medial fronto-orbital gyrus; MOG, middle occipital gyrus; MTG, middle temporal gyrus; PoCG, postcentral gyrus; PrCG, precentral gyrus; SFG, superior frontal gyrus; SOG, superior occipital gyrus; SPG, superior parietal lobule.

We further examined how behavioral manifestations were encoded by the identified hierarchic limbic-sensorimotor-association gradient. As shown in Fig. 5D, the infant rCBF hierarchic distribution regions were largely associated with a specific developmental functioning. In particular, language skill was primarily encoded in sensorimotor cortices, and cognitive was mainly encoded in frontoparietal cortex. Motor skill was encoded in sensorimotor and frontoparietal cortices. These findings suggest that region-specific rCBF increases along the hierarchic limbic-sensorimotor-association gradient are associated with enhanced real-world developmental functioning during infancy.

### Infant rCBF reorganized by spatiotemporally varying energy demand for brain maturation

Given that blood flow delivers glucose and oxygen to different brain regions to meet energy demands, rCBF distributions are tightly associated with the neuronal activities and functional specializations among brain areas ^7^. Consistent with the literature ^7,8^, and shown in Fig. 6A, rCBF is a high-fidelity surrogate measure of cerebral metabolic rate of glucose (CMRglc) measured by FDG-PET. Despite individual differences, a strong spatial association between adult rCBF and CMRglc maps (*r* = 0.594, *P_perm_* < 10^-4^; Pearson correlation with permutation test) was observed (Fig. 6A). However, FDG-PET is usually not applicable to typically developing infants given radiation concerns. From Fig. 6B, infant rCBF is progressively reorganized during development from significantly different distribution at a younger age (e.g. 1.5 months) indicated by lower r value (*r* = -0.035, *P_perm_* < 10^-4^) to a distribution highly aligned with the adult metabolic distribution map at an older age (e.g. 28 months) indicated by higher r value (*r* = 0.635, *P_perm_* < 10^-4^). A reorganization plateau is reached at around 10 months (Fig. 6B), coinciding with the break-point age of global CBF increase (Fig. 1). These findings suggest that the cerebral cortex undergoes a dynamic course of metabolic maturation during infancy, and the hierarchy-dependent age-related changes of rCBF are impacted by the allocation of brain energy demand for brain maturation.

**Figure 6:**
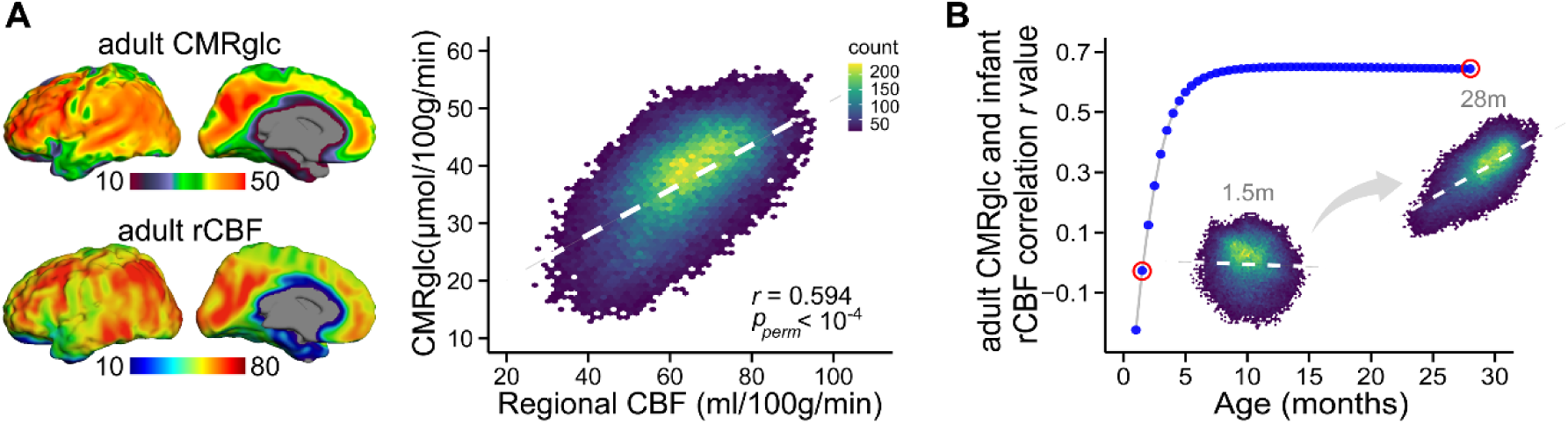
Infant rCBF reorganized by spatiotemporally varying energy demand for brain maturation. **(A)** The *upper-left* panel shows the adult cerebral metabolic rate of glucose (CMRglc) map, acquired from published average maps across 28 healthy adults from Shokri-Kojori et al. (2019). The adult CMRglc map highlights frontal lobe and precuneus as the most metabolically demanding regions. The *bottom-left* panel shows the rCBF map acquired from a healthy adult using the same scanner and identical pCASL protocol as the infant cohort. Shown in the *right* panel, adult rCBF significantly correlated with the adult CMRglc (*r* = 0.594, *P_perm_* < 10^-4^; Pearson correlation with permutation test). **(B)** Spatial distribution of infant rCBF map is progressively reorganized during development and gradually aligns with the adult CMRglc distribution pattern. Spatial alignment between fitted infant rCBF map from 1 to 28 months (m) and adult CMRglc slowly increases, reaching a plateau at 10m of age. RCBF-CMRglc associations at two representative ages (red circled) of 1.5 and 28 months are also shown. Unlikely the weak association between 1.5m infant rCBF and adult CMRglc maps, the distribution of 28m infant rCBF is highly aligned with the adult metabolic distribution map.

## Discussion

We found the emergence of the limbic-sensorimotor-association cortical hierarchy in the infant brain using noninvasive physiological imaging based on cerebral blood flow. This physiological cortical hierarchy reflects changes in regional neural development that likely underlies cortical functional specialization. We also discovered the physiological basis of behavioral manifestation by associating increases in cerebral blood flow in specific cortical regions and corresponding behavioral measures. Our established normative charts of rCBF at the highest resolution available during infancy not only advances current understanding of normal brain development but also provides an anchor point for detecting abnormal brain perfusion (e.g. brain injury) during this vulnerable period. Taken together, we filled the knowledge gap by elucidating the rCBF dynamics underpinning emergence of the critical hierarchical limbic-sensorimotor-association cortical gradient and revealing rCBF-behavior association in infancy through acquiring the highest-resolution infant rCBF available to date.

Maintaining proper CBF dynamics has a profound impact on brain development and function. CBF delivers glucose and oxygen, the main substrates for energy metabolism in the brain. It is closely coupled with the cerebral metabolic rates for glucose (CMRGlu) and oxygen (CMRO2) ^7,8^, the direct measures of the rate of energy consumption. Loss of CBF is accompanied by irreversible damage to neurons and synapses ^41^. The massive and rapid development renders infancy vulnerable to disorders^9,10^. Thus, infant brain is particularly susceptible to cerebral ischemia and stroke, which disrupts CBF, leaving the brain with inadequate energy to support its underlying maturational processes. The developmental pattern of global CBF during infancy increases rapidly and logarithmically with age and coincides with known dynamic changes in brain metabolism ^23,24^, reflecting a phase of escalating metabolic demands associated with the overproliferation of energetically costly synapses and dendritic spines ^42^. Dramatic cellular processes in early developmental brains, such as extensive synaptogenesis, dendritic arborization, glial proliferation, and myelination, are critical for the formation and growth of neural circuits^6^, allowing for rapid brain function development. These intensive developmental processes and increased neural activity necessitate higher metabolic demands, promoting angiogenesis for an elevation in CBF to supply more glucose and oxygen. Moreover, we found that global CBF had a similar rise and break point as the adjusted total daily energy expenditure during infancy ^36^, consistent with the notion that much of infant whole-body metabolism is dedicated to brain development^5^.

We further observed that rCBF was heterogeneously distributed across brain regions during infancy. The spatial pattern of infant rCBF resembled that of local brain metabolism at a certain age. For example, the prominent physiological activity in primary sensorimotor and auditory cortical areas, putamen, and thalamus in 0-6-month-old infants is consistent with higher glucose utilization ^23,24^ in these regions than the rest of the brain. In contrast, by 9 to 12 months, infants’ rCBF patterns qualitatively appear like those of adults, with strong physiological activity in frontal and parietal association cortices, matching the changes in their metabolic patterns as well. We also discovered that infant rCBF increased in a spatially heterogenous fashion, resulting in a shift of rCBF distribution patterns as a function of age. These findings align with prior infant ASL research ^e.g., 29–31^, which noted uneven distributed rCBF and its changing patterns but mainly focused on limited brain regions and lacked sufficient coverage of the entire first 2 years of life. The spatiotemporal heterogeneity of infant rCBF dynamics underlies the emergence of cortical gradient.

Most notably, our study provides the first clear view of physiological hierarchy along the limbic-sensorimotor-association gradient throughout infancy. To our knowledge, such cortical axis with limbic component has not been reported so far, although the sensorimotor-association gradient been proposed for cortical maturation at much later developmental stages such as adolescence and adulthood ^e.g. 9^. The identified physiological hierarchy aligns well with the pronounced and qualitatively distinct regional differences in infant functional connectivity ^20–22,30^, neuroanatomy ^2,16,43^, and transcriptomic profile ^44,45^. For instance, limbic and primary sensorimotor functional networks are relatively well established at birth ^20,21^. In contrast, functional networks involving association cortices (e.g., default mode, frontoparietal, and attention networks) are often recognizable in topographically incomplete forms at birth and then grow rapidly during the first two years of life ^22,30^, gradually establishing the limbic-sensorimotor-association functional gradient. Thus, it is not surprising that rCBF rises greatly in these association cortices to meet the extra energy demand for functional network and gradient formation^30^. Infant brain also undergoes remarkable but non-uniform morphological changes, with lateral temporal, parietal, and frontal cortices expanding more quickly than primary sensorimotor and occipital cortices ^16^. Cortical thickness of the medial orbitofrontal cortex from limbic network reaches its peak earliest with respect to all cortical cortex^16^. White matter develops in a similar spatial pattern^2,12^. For example, limbic fibers (e.g. fornix) appear and myelinate earliest, followed by sensorimotor projection fibers (e.g. corticospinal tract), and then association fibers connecting to frontal areas (e.g. inferior fronto-occipital fasciculus)^2,17–19^. These rapid functional and structural brain changes are reliant on the spatiotemporal regulation of the transcriptome with gene expression during infancy showing differences between primary and association cortical areas ^44,45^ and pronounced enrichments of biological processes related to synaptogenesis and myelination that proceed heterochronically across brain regions^42^. The emergence of the limbic-sensorimotor-association gradient comes at substantial metabolic cost. Developmental refinement of rCBF dynamics throughout infancy reflects the reallocation of the brain’s energy resources to promote the establishment of this cortical gradient. This infant cortical hierarchy with limbic component may transition over time, merging into the sensorimotor-association gradient observed in adolescents and adulthood. Our findings converge with the functional, transcriptomic and structural findings in the literature and fill a gap regarding the physiological hierarchical emergence of the human cortex. The CBF-based physiological hierarchy may underlie the emergence of macrostructural, microstructural, and functional hierarchy.

In parallel with the rapid brain changes for establishing cortical hierarchy, infants begin to walk, talk, and develop increasingly complex behavior. We found that increased rCBF was associated with measures of infant behaviors and development in real world setting. As the majority of brain’s energy is devoted to neural signaling processes that maintain brain function ^46^, increased CBF may ensure that the brain meets the energy demands needed for the formation of large-scale functional networks capable of supporting complex behaviors ^30,47^. Cortical regions with significant rCBF-behavior associations exhibited selectivity for different behaviors, suggesting distinct brain circuits facilitating the development of specific behaviors and functioning. For language development, increased rCBF was predominantly seen in sensorimotor regions that contributing to the execution of motor activities. This is in line with the finding of sensorimotor network influences speech and language acquisition in infancy ^48^. In contrast, cognitive development was more associated with rCBF in frontoparietal areas. It is noteworthy that the frontoparietal network plays an important role in infants’ social cognitive development ^49^. Understanding how rCBF dynamics support the emergence of real-world infant behavior and developmental functioning may provide potential biomarkers for early detection of disorders in infants, as adaptive behavior delays during infancy have been linked to later diagnoses of neurodevelopmental conditions such as autism ^50^.

Prior studies that focused on small subsets of brain regions or incomplete coverage of the first 2 postnatal years found varied rCBF distribution and changes in infant brains ^e.g., 29–31^, but our study is the first to delineate the rCBF hierarchy throughout infancy. Low-resolution maps in prior work hampered the capacity to precisely chart the infant rCBF topography. Capitalizing on a cutting-edge 3D multi-shot, stack-of-spiral pCASL with background suppression ^33,34^, the much higher imaging resolution in present study distinguishes rCBF heterogeneity across proximate brain areas. The resultant highest infant rCBF maps to date with 2.4 times or more resolution improvement than previous studies change the paradigm and enable capturing the unprecedented details of the infant whole-brain physiological hierarchy throughout infancy. According to the ASL consensus paper ^27^, 3D acquisition is the recommended ASL readout method because it is SNR-efficient and can be optimally combined with background suppression to effectively reduce physiological noise when compared to 2D readout. In this study, only 3D ASL data from cohort-1 were used for the main results, whereas 2D ASL data from cohort-2 was used for validation only. Furthermore, we mitigated the impact of in-scanner head motion on our results by rigorously assessing image quality for data inclusion and co-varying for motion in all analyses. These procedures limit the likelihood that the reported findings are due to in-scanner head motion. Despite slightly more female in cohort-1, sex was systematically accounted for as a covariate in all analyses to minimize any potential effects. We demonstrated that rCBF is strongly correlated with local brain metabolism at rest despite that rCBF and CMRglc were acquired from different adult individuals. The observed age-related rCBF change pattern in infants of this study is also aligned with CMRglc measures from previous studies ^23, 24^ However, caution needs to be taken for the interpretation as rCBF is not a direct measure of the rate of brain energy consumption. With only cross-sectional data presented in this study, longitudinal studies are warranted to minimize the inter-subject variability when delineating age-related rCBF dynamics in infancy. Future studies can investigate how combining rCBF with neuroimaging measures from other modalities, such as functional MRI and diffusion MRI, might reveal multidimensional principles of infant brain maturation. This work is a step towards the normative rCBF milestone for clinical translation. With rapid MR technical advances in hardware and sequence, the high-resolution rCBF currently obtained with a research protocol can be soon translated into clinical settings. It is recognized that a barrier to ASL clinical translation is the lack of standardization of these perfusion MRI methods across scanners and across sites. With the growing effort on generating consensus ^e.g. 27,51^ and promoting standardization for imaging biomarker (e.g., QIBA: Quantitative Imaging Biomarkers Alliance from the Radiological Society of North America, https://qibawiki.rsna.org), variation of rCBF measurements across vendors and sites will likely be reduced over time as more studies incorporate ASL and more clinical data are collected and validated.

## Acknowledgements

This work was supported by grants from National Institute of Health: R01MH092535, R01MH125333, R01EB031284, R01MH129981, R21MH123930, R01EB031080, R01HD093776, R01MH107506, and P50HD105354.

## Author contributions

M.O. and H.H. designed the research. M.O., J.A.D., J.L.H, K.L.S., E.S.K., J.C.E., Y.P., and H.H. performed the research. M.O., J.A.D. and H.H. contributed new methodology and analytic tools. M.O. and H.H. supervised the collection of neuroimaging data. M.O. analyzed data, and generated figures and tables. M.O. wrote the original draft and all authors (M.O., J.A.D., J.L.H, K.L.S., E.S.K., J.C.E., Y.P., and H.H.) reviewed and revised the final darft.

## Methods

### Participants

As shown in Supplementary Table 1, a total of 134 infants from two independent cohorts were recruited in this study. In cohort-1 (*N* = 78), we collected high-resolution pCASL of isotropic 2.5 mm and PC perfusion MRI in infants at the Children’s Hospital of Philadelphia. In cohort-2 (*N* = 56), infants underwent pCASL and PC perfusion MRI at the Beijing Children’s Hospital. We combined PC-MRI data from two cohorts to investigate the developmental model of global CBF in a large sample (Fig. 1; *N* = 119; 63 from cohort-1 and 56 from cohort-2). No cohort effects were observed (Supplementary Table 2). We used high-resolution pCASL data from cohort-1 (*N* = 76) to delineate the developmental models of regional CBF during infancy (Fig. 2-6). For validation, key findings reported for voxel-wise age effects on regional CBF during infancy were replicated using the pCASL data collected from infant cohort-2 (*N* = 48; Supplementary Fig. 5). Below are detailed descriptions of each infant cohort.

Cohort-1 is part of an ongoing study investigating typical brain development during infancy at the Children’s Hospital of Philadelphia, Philadelphia, USA. Participants were recruited from the greater Philadelphia metropolitan area with assistance from the Clinical Reporting Unit at Children’s Hospital of Philadelphia for identifying eligible participants and through study flyers. All recruited infants were enrolled as typically developmental infants (i.e. brain imaging not clinically indicated). They were recruited completely for research purposes, which involved studying normal infant brain development. All infants were selected after rigorous screening procedures conducted by an experienced psychologist (ESK) based on their medical history and family medical history. Exclusionary criteria for infants were: premature birth or significant birth complications; known neurological disorders; genetic conditions with known neurological or neurodevelopmental sequelae; severe tics, or severe head trauma; known developmental or medical diagnoses; major congenital heart disease or congenital infection; uncorrectable sensory (hearing, visual) issues; having a sibling or parent with a diagnosis of autism spectrum disorder or schizophrenia; and 1.5 or more standard deviations below the mean on the behavioral assessment of developmental milestones. From the 275 infants who were originally recruited to a larger normal brain development study, 78 infants who completed scans of either PC-MRI or pCASL perfusion MRI participated in this study as cohort-1. Out of 78 infants, advanced pCASL was successfully acquired from 76 infants aged 1.2 to 28 months (mean/standard deviation 10.98 ± 6.82 months; 32M/44F), and PC-MRI was acquired from 63 infants aged 1.5 to 28 months (mean/standard deviation 9.84 ± 7.36 months; 26M/37F). All procedures for cohort-1 were approved by the Institutional Review Board at the Children’s Hospital of Philadelphia in compliance with ethical regulations and standards (Approval number IRB16-013203). Written informed consent was obtained from all infants’ parents.

Cohort-2 included 56 infants aged 1.4 to 27.7 months (mean/standard deviation 14.49 ± 6.83 months; 34M/22F) and recruited at Beijing Children’s Hospital, Beijing, China. These infants were referred to MR imaging due to simple febrile convulsion (*N* = 19), convulsion (*N* = 21), diarrhea (*N* = 13), or sexual precocity (*N* = 3). All infants had normal neurological examinations documented in their medical records. Exclusion criteria include known nervous system diseases, or a history of neurodevelopmental, psychiatric, or systemic illness. The clinical history of each infant was carefully inspected to rule out developmental abnormalities. No brain abnormalities were detected on the MRI scans of these infants, based on the reading of an experienced pediatric radiologist (YP). All infants from this cohort-2 had PC-MRI acquired, and 48 of them had pCASL acquired. All procedures for cohort-2 were approved by Beijing Children’s Hospital Research Ethics Committee (Approval number 2016-36) and every participant’s parents provided signed consent.

### Multimodal MR image acquisition

In general, multimodal brain MRIs for infants were obtained at higher resolutions in cohort-1 than in cohort-2. Brain scans of infants in cohort-1were collected during natural sleep on a 3T Siemens Prisma scanner with a 32-channel head coil at the Children’s Hospital of Philadelphia. Brain scans of infants in cohort-2 were collected on a 3T Philips Achieva scanner with an 8-channel head coil at the Beijing Children’s Hospital. Infants from cohort-2 were under sedation with orally administered chloral hydrate at a dose of 0.5 ml/kg and no more than 10 ml in total during scanning. Previous research^52, 53^ suggested no significant effect of chloral hydrae on cerebral blood flow or sensory function. Earplug and headphones were used to reduce noise exposure. Below are detailed parameters for each MRI modality.

#### Structural MRI

Prior to perfusion MRI acquisition, structural MRIs were acquired. In cohort-1, high-resolution structural MRIs were acquired with magnetization-prepared, rapid acquisition gradient-echo T1-weighted (MPRAGE, T1w) images (TR = 2400ms; TE = 2.24ms; TI = 1060ms; flip angle = 8°; sagittal FOV = 256×256mm^2^, matrix = 320×320, slice number = 208; resolution = isotropic 0.8mm, scan duration = 6mins38s) and a variable-flip-angle turbo-spin-echo T2-weighted (Siemens SPACE, T2w) images (TR = 3200ms; TE = 564ms; turbo factor = 314; sagittal FOV = 256×240mm^2^, matrix = 320×300, slice number = 208; resolution = isotropic 0.8mm, scan duration = 5mins57s). In cohort-2, structural MRI was acquired with T1-weighted images (TR = 8.28ms; TE = 3.82ms; TI = 1100ms; flip angle = 12°; sagittal FOV = 200×200mm^2^, matrix = 200×200, slice number = 150; resolution = isotropic 1mm, scan duration = 3.7mins).

#### Perfusion MRI

We collected pCASL and PC-MRI in both cohorts to measure regional and global CBF, respectively. To position the PC-MRI imaging planes, we also acquired angiogram. MR images from a representative infant were used to demonstrate pCASL, PC-MR and angiogram acquisitions (Supplementary Fig. 1A).

In cohort-1, all high-resolution pCASL scans were acquired with a 3D multi-shot, stack-of-spirals pCASL sequence ^33,34^ with the following parameters: TR = 4000ms; TE = 9.4ms; four-shot acquisition; FOV = 192×192mm^2^, matrix = 76×76; 48 slices; slice thickness/gap = 2.5/0mm, effective voxel resolution = 2.53×2.53×2.5mm^3^; labeling duration=1600ms; post labeling delay (PLD) = 1800ms; number of controls/labels = 10 pairs; center of labeling slab located between cervical vertebrae C2 and C3 (Supplementary Fig. 1A); background suppression to suppress approximately 90% of the signal contributed from static tissue to reduce physiological noise, and scan duration = 7mins6s. For each pCASL scan, a pair of M_0_ images were acquired and averaged for rCBF quantification with longer TR, no magnetization preparation, and the same readout scheme ^33^. A time-of-flight (TOF) angiogram was acquired with axial slices encompassing a slab covering the foramen magnum. The imaging parameters of the angiogram were: TR = 21ms; TE = 3.42ms; flip angle = 18°, FOV = 200×180mm^2^, matrix = 256×144; 30 slices; slice thickness/gap = 2/0mm, effective voxel resolution = 0.78×0.78×2.0mm^3^; thickness of saturation slab above the imaging slab = 40mm, and scan duration = 47s. The four feeding arteries, including bilateral internal carotid artery (ICA) and vertebral artery (VA), could be well visualized in the three-dimensional reconstructed angiogram on the middle and right panels of Supplementary Fig. 1A. Based on the angiogram, the imaging planes for PC-MRI of ICAs were placed at the level of the foramen magnum and the imaging planes for PC-MRI of VAs were placed between the two turns in V3 segments (at approximately the level of the C1 vertebral column) (Supplementary Fig. 1A). Image parameters of PC-MRI were: TR = 23.2ms; TE = 5.76ms; flip angle = 18°; FOV=200×150mm^2^, matrix = 256×120; 14 slices; slice thickness/gap = 2/0mm, effective voxel resolution = 0.78×0.78×2.0mm^3^; maximum velocity encoding = 40 cm/s; non-gated; 2 repetitions, and scan duration of each artery = 33s. As shown in the right panels of Supplementary Fig. 1A, the cross-section of the target artery with higher intensity can be observed in each PC image.

In cohort-2, infant pCASL images were acquired with a 2D multi-slice echo-planar imaging sequence with following parameters: TR = 4100ms; TE = 15ms; FOV = 230×230mm^2^, matrix = 84×84; 20 slices; slice thickness/gap = 5/0mm, effective voxel resolution = 2.74×2.74×5 mm^3^; labeling duration=1650ms; PLD = 1600ms; number of controls/labels = 30 pairs; center of labeling slab located at the junction of spinal cord and medulla (65 mm below central slab of imaging plane), and scan duration = 4.2mins. An auxiliary scan with identical readout module to pCASL but without labeling was acquired for estimating the value of equilibrium magnetization of brain tissue. Similarly, a TOF angiogram was acquired with following parameters: TR = 20ms; TE = 3.45ms; flip angle = 30°, FOV = 100×100mm^2^, matrix = 100×100; 30 slices; slice thickness/gap = 1/0mm, effective voxel resolution = 1×1×1mm^3^, and scan duration = 28s. PC-MRI were acquired with following parameters: TR = 20ms; TE = 10.6ms; flip angle = 15°; FOV = 120×120 mm^2^; matrix size = 200×200; single slice, effective voxel resolution = 0.6×0.6×3mm^3^; maximum velocity encoding= 40cm/s; non-gated; 4 repetitions, and scan duration of each artery = 24s. More details for cohort-2 can be found in our previous work^30^.

#### Infant MRI scan preparation for cohort-1

Infant MRI has been known to be challenging. We developed specific preparations to successfully acquire multimodal MR imaging from infants during natural sleep. Before scheduling the visit, a telephone screening is completed with the child’s caregiver to collect information, including infant’s typical nap and feeding schedule, and sleeping preferences (i.e., use of pacifier, white noise, swaddling, and crib/napping environment). The MRI scan is usually scheduled at the infant’s nap or bedtime. After scheduling, we send a confirmation email that outlines MRI visit details, what to expect, what to bring, and audio files of MRI noises to practice playing for their infant during nap time at home. Both infant and caregiver are required to change into MRI-safe hospital gowns during the imaging visit, and warmed blankets are provided for infant to be swaddled in. The MRI waiting room is used for caregivers to begin preparing for their infants to go down for their nap. Caregivers are encouraged to follow their infants’ typical nap and bedtime routines as closely as they can (i.e., diaper change, swaddle, reading a book, singing). When they are ready to enter the MRI room, the research assistant and technologist escort them into the scanning room after being screened for metal in or on the body. All lights are turned off and white noise or other soothing sounds are played inside the scanning room at the parent’s request. An MRI-compatible rocking chair is set up for the caregiver to feed and rock the infant to sleep in their arms. If rocking the infant in their arms is not possible, we allow the caregiver to let the infant fall asleep on an MRI safe bed before transitioning them to the MRI scanner table. At this time, MiniMuffs for ear protection to reduce scanner noise are placed on the infant’s ears. The caregivers are encouraged to wait 5-10 minutes after their infant initially falls asleep to ensure they are in a deeper sleep for transferring onto the MRI scanner table. Once asleep, the caregiver assists in gently placing the infant on the scanner table. To minimize motion during the scan, the MRI technologist positions the infant’s head in the coil using foam pads and cushions around the head.

### Infant behavior and real-world developmental functioning

Only infants from cohort-1 had their behavior and developmental functioning assessed. To examine the associations of regional CBF dynamics with measures of infant behavior, we analyzed the Bayley scales of infant and toddler development ^54^ that was completed during infants’ imaging visit at the Children’s Hospital of Philadelphia. This neurodevelopmental assessment was conducted by a certified neurodevelopmental psychologist (ESK), who was blinded to the infant MR findings. Fifty-three of the seventy-six infants who had high-resolution pCASL acquired in cohort-1 completed the Bayley scale to assess their behavior and real-world developmental functioning in three domains: motor, language, and cognitive. Specifically, the cognitive scale estimates general cognitive functioning on the basis of nonverbal activities; the language scale estimates receptive communication as well as expressive communication including the ability to communicate through words and gestures; and the motor scale estimates both fine motor and gross motor. The Bayley scale is age standardized and widely used in both research and clinical settings. The three Bayley domains’ composite scores are expressed as standard scores with a mean of 100 and a standard deviation of 15, with higher scores indicating better performance. Unlike cognitive, language and motor scales reliably obtained using items administered to the infant by a certified neurodevelopmental psychologist, other two scales from Bayley (social-emotional and adaptive scales) obtained from primary caregiver heterogeneous responses to questionnaires were not included in this study.

### Multimodal MR imaging processing with estimated motion

We estimated global and regional CBF from PC and pCASL perfusion MRI for infant datasets from both cohorts using the preprocessing methodologies described below.

#### Global CBF estimation with PC perfusion MRI

PC-MRI provides a quantitative measurement of the flow velocity of a given blood vessel. By integrating over the cross-section of the vessel, blood velocity can be converted to flow rate. The flow rates from four feeding arterials can then be used to calculate global CBF ^35^ and formulated as

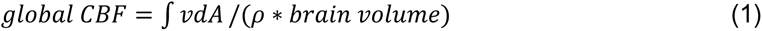

where *v* is the blood flow velocity in the ICAs and Vas; *A* is the cross-sectional area of blood vessel with a unit of mm^2^; and ρ is the brain tissue density assumed as 1.06 g/mL^55,56^. Brain volume was measured from structural MRI as parenchyma volume (gray matter + white matter volume).

#### Regional CBF estimation with pCASL perfusion MRI

After head motion correction of the pCASL perfusion MRI, we estimated rCBF using the protocol similar to that in our previous publication^28^. Briefly, rCBF was measured using a model described in the ASL consensus paper ^27^:

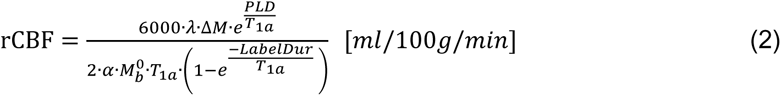

where *ΔM* is the dynamic-averaged signal intensity difference between in the control and label images; *λ* is the blood-brain partition coefficient in ml/g (0.9 ml/g ^56^); PLD is the post labeling delay time; LabelDur is the labeling duration; *α*, the labeling efficiency, is 0.86 predicted by the fitting between labeling efficiency and blood velocity in the previous study ^57^; T_1a_ is T_1_ of arterial blood (1800 ms ^58,59^). The value of equilibrium magnetization of brain tissue (*M*^0^) was obtained from the M_0_ images from the acquisition. The labeling efficiency α and T_1a_ could vary considerably across participants, especially in infants. Thus, we used global CBF from PC-MRI to calibrate rCBF measures from pCASL, as described previously ^28,30,57^. Specifically, rCBF was calibrated by applying the scalar factor making averaged rCBF equal to global CBF. Previous validation studies have shown that this procedure enhances accuracy and reliability of rCBF quantification ^57^. The pCASL imaging parameters for infants, such as PLD, were selected in the interval between imaging parameters of neonates and those of children, with a PLD of 2000ms for neonates and 1500ms for children recommended by the ASL consensus paper ^27^. Previous studies reported the mean arterial transit time (ATT) in infants less than 1500ms ^59^. Following the consensus paper ^27^ suggestion that the PLD should be larger than the ATT, a tailored PLD of 1600-1800ms which is larger than ATT of infants was selected for the infant cohorts to minimize the effect of PLD on rCBF across subjects and brain regions.

We scanned an infant (18 months old) twice to demonstrate the reproducibility of the infant pCASL protocol in cohort-1 (see *Multimodal MR imaging acquisition*). To quantify the test-retest reproducibility of rCBF measurement, we further calculated the intraclass correlation coefficient (ICC) with rCBF maps estimated from these two pCASL scans (Supplementary Fig. 1B-C).

#### Motion metric for pCASL perfusion MRI

To quantify the in-scanner motion of pCASL, a widely used measurement of mean relative displacement (MRD) across image volumes from previous ASL and functional MRI studies ^e.g. 60,61^ was adopted. We used the same procedures from previous studies ^60,61^ to obtain the MRD measurement. First, all pCASL images were motion-corrected using the MCFLIRT function from the FSL package ^62^. The re-alignment parameters, six motion parameters consisting of three translations and three rotations, estimated from MCFLIRT across pCASL volumes, were condensed to a single vector representing the root mean squared (RMS) volume-to-volume displacement of all brain voxels. This one-dimensional motion timeseries were further used to calculate the RMS displacement relative to the preceding volume (i.e., relative RMS displacement). In order to provide a summary measure of in-scanner motion for each subject, the MRD measurement was estimated as the mean value of the relative RMS displacement vector. Subjects or image volumes with excessive motion (MRD or relative RMS displacement > 0.5mm) were excluded ^60^. MRD values from all 76 infants (0.22 ± 0.13mm) were less than 0.5mm (Supplementary Table 3). Infant’s in-scanner motion was also included as a covariate in the subsequent analyses of developmental models of rCBF and the association analyses with infant behavior. In addition to the above-mentioned procedures that mitigate potential motion effects, high-resolution rCBF maps in cohort-1 were acquired with 3D multi-shot, stack-of-spiral pCASL with background suppression, a recommended ASL protocol by the ASL consensus paper ^27^. The protocol improves robustness against motion artifacts through stack-of-spiral acquisition for high oversampling in the center of k-space at every shot, and provides high SNR for detecting the ASL signal for rCBF calculation with the background-suppressed 3D readout ^27, 63^.

### Group-averaged rCBF maps generation

To generate the group-averaged rCBF maps across infancy, we registered rCBF maps from all infants in cohort-1 into the JHU brain template in the Montreal Neurologic Institute (MNI) space ^64^ (2mm isotropic resolution with 90×109×90 matrix) utilizing the contrasts of T1w images to drive the registration. Briefly, infants’ T1w images were first co-registered to the M_0_ image from pCASL perfusion MRI in the native rCBF space. Then, a 12-parameter affine registration was used to transform the co-registered T1w image of each infant to the T1w image in the template space, followed by a non-linear transformation (LDDMM: large deformation diffeomorphic metric mapping). By applying the same individual transformations, infant’s rCBF maps were then projected to the template space. The registration procedures were conducted using DiffeoMap software (www.mristudio.org). We divided the 76 infants from cohort-1 into six age groups (i.e., 0-3, 3-6, 6-9, 9-12, 12-18, and 18-28 months; Supplementary Table 3) to characterize the dynamic rCBF changes during infancy. The corresponding rCBF maps from each age group were then averaged in template space to generate the group-averaged maps. For each infant, rCBF values from the 144,237 gray matter voxels as defined by the JHU brain atlas ^64^ were used for subsequent analyses of voxel-wise developmental models of rCBF.

### Delineate the developmental models of global and regional CBF

#### Identify the developmental model for age-related change of global CBF

To quantify the developmental curve of global CBF during infancy, we first identified the best model for the relationship between global CBF and age. Model selection for developmental curves of global CBF utilized the Akaike information criterion (AIC) and Bayesian information criterion (BIC) to quantify the relative goodness of fit among different candidate models^65,66^, including linear, exponential, logarithmic, Poisson, and quadratic polynomial for fitting the age-related curves of global CBF during infancy (Supplementary Fig. 2). The AIC = -2ln(L) + 2k, where L = maximum likelihood of model and k = number of parameters. Since the absolute AIC value is highly specific to the dataset, the relative AIC (ΔAIC_i_ = AIC_i_ – AIC_min_) are more widely used to rank models. The value of the ΔAIC_i_ of the relatively best model is set to 0. The likelihood of a model can be estimated by *e*^−1/(2ΔAICi)^. The likelihood of each model divided by the sum of the likelihood for all models to estimate the AIC weight (wAIC), which can be interpreted as the probability of the model to be the best model among the candidate models (Supplementary Table 2). All analyses were conducted in R version 4.1.1 ^67^. The AIC results showed that the logarithmic model had more than 99% probability to be the best model for characterizing global CBF development during (Supplementary Table 2). We also investigated the effects of sex and cohort on developmental curve of global CBF with a multivariate regression model (e.g., Eq. 3).

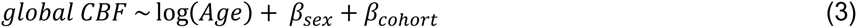

#### Identify the developmental models for age-related change of regional CBF

After analyzing the effect of age on global CBF, we sought to evaluate the spatial distribution of age effects on rCBF. Considering the logarithmic model best fitted the global CBF growth during the same developmental stage, we adopted the same model here for voxel-wise developmental models of rCBF. Infant sex and in-scanner head motion were included as covariates with each logarithmic model. Head motion was quantified as MRD (see *Motion metric for pCASL perfusion MRI*). This model can be summarized using the formula in Eq. 4:

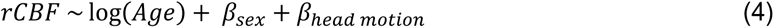

We used Bonferroni correction to further control the type 1 error in the age effects quantification on rCBF (*Z* map). Given a sample of 76 subjects (cohort-1) and age effects across the 144,237 gray matter voxels, corrected *P* < 0.05 corresponds to uncorrected *P* < 3.47×10^-7^ or *Z* > 5.1. To visualize the dynamic distribution patterns of rCBF throughout infancy, we constructed fitted rCBF maps at specific ages ranging from 1 to 28 months with an increment of 0.5 months based on the best-fit development curves across cortex (Supplementary Fig. 6-7). In estimating the rCBF at each age, we assumed the head motion as 0.22mm (i.e., the averaged MRD value from all subjects) and averaged the sex effect from both genders. The resulting rCBF maps were then animated to create a quantitative time-lapse movie of spatiotemporal changes of rCBF during infancy (Supplementary Movie 1).

For validation, the voxel-wise age effects on rCBF (Z map) during infancy were also replicated using the infant perfusion MRI from cohort-2. In order to evaluate the alignment of age effects on rCBF from two cohorts, Pearson correlation between two Z maps was calculated (*r_oberserved_*). The significance of the alignment was further quantified using a nonparametric permutation test. Specifically, surrogate Z maps (*N* = 10,000) were generated by re-assigning voxel-wise values from Z map of cohort-2. A distribution of correlation coefficients was constructed by correlating these surrogate maps with Z map from cohort-1. The proportion of 10,000 null correlations (*r_null_*) that exceed the observed correlation coefficient between the two Z maps (*r_oberserved_*) is defined as an empirical *P_perm_* value. *P_perm_* < 0.05 indicates significant alignment of age effects on rCBF from two infant cohorts (Supplementary Fig. 5).

#### Segmented regression analyses of infant CBF

To identify distinct phases of CBF changes during infancy, we employed the segmented regression approach, which has also been used in previous studies ^e.g., 11, 36^ to detect break points between different phases in age-related curves of imaging measures. We used segmented regression analysis to determine the break points in the relationship between infant CBF (i.e., global and regional) and age with the *Segmented* (version 1.4-1) package ^68^ in R version 4.1.1. The approach requires the user to specify the number of breakpoints, and then identifies each break-point location or value. We examined a range of models with 0 to 3 break points in the *segmented* function for infant CBF. Increase in the number of breakpoints from 0 to 1 improved the model adjusted R^2^ and standard error. Statistical significance was assessed using analysis of variance (ANOVA) to compare the models (e.g., linear and biphasic linear models). Increasing the number of breakpoints to 2 or 3 did not significantly improve the model. We therefore selected the biphasic model as the best fit for infant CBF in this dataset. A similar one break-point segmented regression approach was conducted for global CBF, and regional CBF from clusters identified by below clustering analyses. Segmented regression results are shown in Supplementary Table 4.

### Clustering analyses of infant rCBF

To reveal the spatiotemporal heterogeneity of rCBF change in the infant brain, we parcellated brain regions using a data-driven clustering approach named non-negative matrix factorization (NMF), a reliable parcellation technique that has been widely used in neuroimaging studies ^e.g^ ^39,69^. Instead of employing established cortical parcellations based on adult brains, this approach allows us to discover the infant-specific cortical topography of rCBF developmental regionalization. The NMF method generates a parts-based representation of all infants’ rCBF maps by grouping cortical voxels that change in a similar fashion, thus facilitating interpretation of rCBF developmental regionalization during infancy. Specifically, NMF factorizes a tall non-negative data matrix ***X*** ∈ *R*^*M*×*N*^ into two non-negative matrices: a component matrix ***W*** ∈ *R*^*M*×*K*^ and a coefficient matrix ***H*** ∈ *R*^*K*×*N*^. In other words, the new variables ***W***, termed hereafter components or clusters, were constructed as a linear, non-negative combination of the original variables. NMF aggregated variance in the components by positively weighting original variables (i.e., rCBF) that tend to co-vary. The scalar *K* represents the number of components (i.e., clusters), which is usually small (i.e., *K* ≪ *M* and *K* ≪ *N*). In contrast to clustering approaches that assign each cortical voxel to a single component, NMF yields a soft (probabilistic) parcellation. This soft parcellation can be converted into hard (discrete) regionalization by assigning each cortical voxel to a specific component according to its highest weight. Estimating the components with NMF can be formulated as:

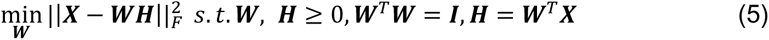

In this study, ***X*** ∈ *R*^*M*×*N*^ is a large non-negative data matrix containing rCBF values from all infants registered to the template space, where *M* and *N* represent the number of cortical voxels and subjects, respectively. To find an appropriate number of components (i.e., *K*) for rCBF growth, we adopted Davies-Bouldin criterion ^70^ and Calinski-Harabasz criterion ^71^ to evaluate the performance of clustering analyses. The Davies-Bouldin index represents average similarity measure of each cluster to its most similar cluster, and a lower value of the index means the clusters are well separated. The Calinski-Harabasz index represents the ratio of the sum between-cluster dispersion and within-cluster dispersion for all clusters, and higher value of the index means the clusters are dense and well separated. We selected component number of 3 with the best criteria scores for categorizing rCBF development, after testing the clustering analyses with *K* ranging from 2 to 20 (Supplementary Fig. 8). The components can be represented as soft cluster assignment maps (Fig. 4A with *K* = 3), where the distribution of the coefficient values of rCBF exhibiting similar statistical properties peaks for the same component.

### Associations of infant rCBF dynamics with infant behavior

We evaluated associations between infant rCBF and real-world developmental functioning at each cortical voxel using generalized additive models (GAMs) with penalized splines to account for linear and nonlinear effect of age ^40^. Infant sex, in-scanner head motion, and family socioeconomic status (SES) were included as covariates in the model (Eq. 6). Here, infant real-world developmental functioning was quantified with Bayley scale across three domains: motor, language, and cognitive (see *Infant behavior and real-world developmental functioning* for details). Family SES was derived from parental education and occupation using the Amherst modification of the Hollingshead two-factor index^72^. Among the infants who had high-resolution pCASL acquired in cohort-1, only 49 of them had both Bayley scale and family SES information completed and data from these 49 infants were included in the association analyses. Cohort-2 was not included in the rCBF-behavior analyses. We used the *mgcv* (version 1.8-39) package in R version 4.1.1 for the analyses.

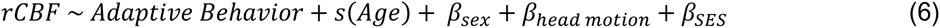

To avoid false positive results, the threshold for voxel-wise across-subject correlation maps was set to a corrected *P* < 0.05, which corresponded to an uncorrected single voxel significance level of *P* < 0.05 (*t* > 2.02) and a minimum cluster extent of 100 continuous voxels (800 mm^3^). The cortical region to which each cluster belongs was identified with infant-specific NMF components (see *Clustering analyses of infant rCBF* for details) and JHU brain atlas ^62^ (Supplementary Table 5). The rCBF values of these cluster voxels were measured to calculate the average rCBF at the identified clusters for each infant. Subsequently, cluster-level across-subject association analyses were performed on the averaged rCBF values and 3 Bayley scores using Eq. 6. Familywise error-corrected significance threshold at cluster level was set at 0.05/3 = 0.0167. Association between cluster-level averaged rCBF and infant developmental functioning was also conducted while controlling for covariates (i.e., age, sex, in-scanner head motion, and family SES).

### Associations of infant rCBF topography with cerebral metabolism in adult brain

To assess the spatial alignment between rCBF and cerebral metabolism, we conducted a Pearson correlation analysis between the rCBF maps and the cerebral metabolism map in the adult brain. To assess the developmental reorganization of infant rCBF, we further conducted Pearson correlation analyses between the fitted rCBF maps across infancy (see *Identify the developmental models for age-related change of regional CBF* for details) and the cerebral metabolism map in adult brain. Cerebral metabolism rate of glucose consumption (CMRglc) with a unit of µmol/100g/min, estimated by ^18^F-flurodeoxyglucose; fluorodeoxyglucose-positron emission tomography (FDG-PET), was used to quantify the cerebral metabolism. The group-averaged (*N* = 28) CMRglc map for adult brain was obtained from the publicly available dataset (see details in reference ^73^; https://github.com/eshoko/COMET) and was registered to the template of infant rCBF maps using linear and nonlinear registrations from FSL package. Association analyses were then performed between the fitted rCBF maps and registered CMRglc map. Permutation tests with 10,000 surrogate maps were used to estimate the significance of these alignments across infancy. Details on permutation testing can be found in the above section on developmental models for age-related change of regional CBF, which describes the alignment of age effects on rCBF across two cohorts.

